# Transcription near arrested DNA replication forks triggers ribosomal DNA copy number changes

**DOI:** 10.1101/2023.12.21.572944

**Authors:** Mariko Sasaki, Takehiko Kobayashi

**Affiliations:** Laboratory of Gene Quantity Biology, Center for Frontier Research, National Institute of Genetics, 1111 Yata, Mishima, Shizuoka 411-8540, Japan; The Graduate University for Advanced Studies, SOKENDAI, 1111 Yata, Mishima, Shizuoka 411-8540, Japan; Laboratory of Genome Regeneration, Institute for Quantitative Biosciences, The University of Tokyo, 1-1-1 Yayoi, Bunkyo-ku, Tokyo 113-0032, Japan; Department of Biological Sciences, Graduate School of Science, University of Tokyo, 1-1-1 Yayoi, Bunkyo-ku, Tokyo 113-0032, Japan; Collaborative Research Institute for Innovative Microbiology, The University of Tokyo, 1-1-1 Yayoi, Bunkyo-ku, Tokyo 113-0032, Japan

**Keywords:** ribosomal RNA gene (rDNA), DNA copy number changes, extrachromosomal circular DNA, chromosome rearrangements, noncoding RNA transcription, Sir2, replication fork arrest, DNA double-strand breaks, homologous recombination, genome stability

## Abstract

DNA copy number changes as a consequence of chromosomal structural rearrangements or the production of extrachromosomal circular DNA. Here, we demonstrate that the histone deacetylase Sir2 maintains the copy number of budding yeast ribosomal RNA gene (rDNA) by suppressing the initiation of homologous recombination (HR)-mediated repair of DNA double-strand breaks (DSBs) that are formed upon DNA replication fork arrest in the rDNA. Sir2 represses the transcription from the regulatory promoter E-pro, which is located near the fork arresting site. When Sir2 is absent, transcription is stimulated by E-pro but terminated by arrested replication forks. This transcription–replication collision enhances DSB formation and induces DSB end resection and HR-mediated repair that is prone to chromosomal rDNA copy number changes and the production of extrachromosomal rDNA circles. Therefore, repression of transcription near arrested replication forks is critical for the maintenance of genome stability by directing DSB repair into the HR-independent, rearrangement-free pathway.

## INTRODUCTION

Changes in DNA copy number are introduced by deletion and/or amplification of a DNA segment on the chromosome. DNA copy number changes can alter gene expression levels and compromise cellular integrity, driving tumorigenesis and numerous other diseases [reviewed by (Sasaki et al., 2010; Carvalho and Lupski, 2016; Harel and Lupski, 2018)]. DNA copy number can also increase via extrachromosomal circular DNAs that are derived from the chromosome (Verhaak et al., 2019). Nearly half of cancer cells contain extrachromosomal circular DNAs (ecDNAs) that carry proto-oncogenes, which are referred to as double minutes or ecDNAs. The accumulation of ecDNAs leads to an increase in the copy number and expression levels of proto-oncogenes, driving tumorigenesis. Thus, understanding the molecular mechanisms underlying DNA copy number changes that occur not only on chromosomes but also via extrachromosomal circular DNAs is essential.

Ribosomal DNA (rDNA) is essential for producing ribosomal RNAs (rRNAs). The budding yeast genome contains ∼150 rDNA copies, which are tandemly arrayed at a single locus on Chr XII (Fig. 1A). Each copy contains 35S and 5S rRNA transcription units, which are separated by intergenic spacers (IGSs). IGS1 contains a replication fork barrier (RFB) sequence and a bidirectional RNA polymerase II (RNAP II) promoter, E-pro, which synthesizes noncoding RNA. IGS2 contains an origin of DNA replication and a cohesin-associated region (CAR). When the rDNA copy number is reduced by less than half, cells induce expansion of the rDNA array (Kobayashi et al., 1998; Kobayashi et al., 2001; Sasaki and Kobayashi, 2023). However, once rDNA copies are restored to the normal level, further expansion is suppressed.

**Figure 1.**
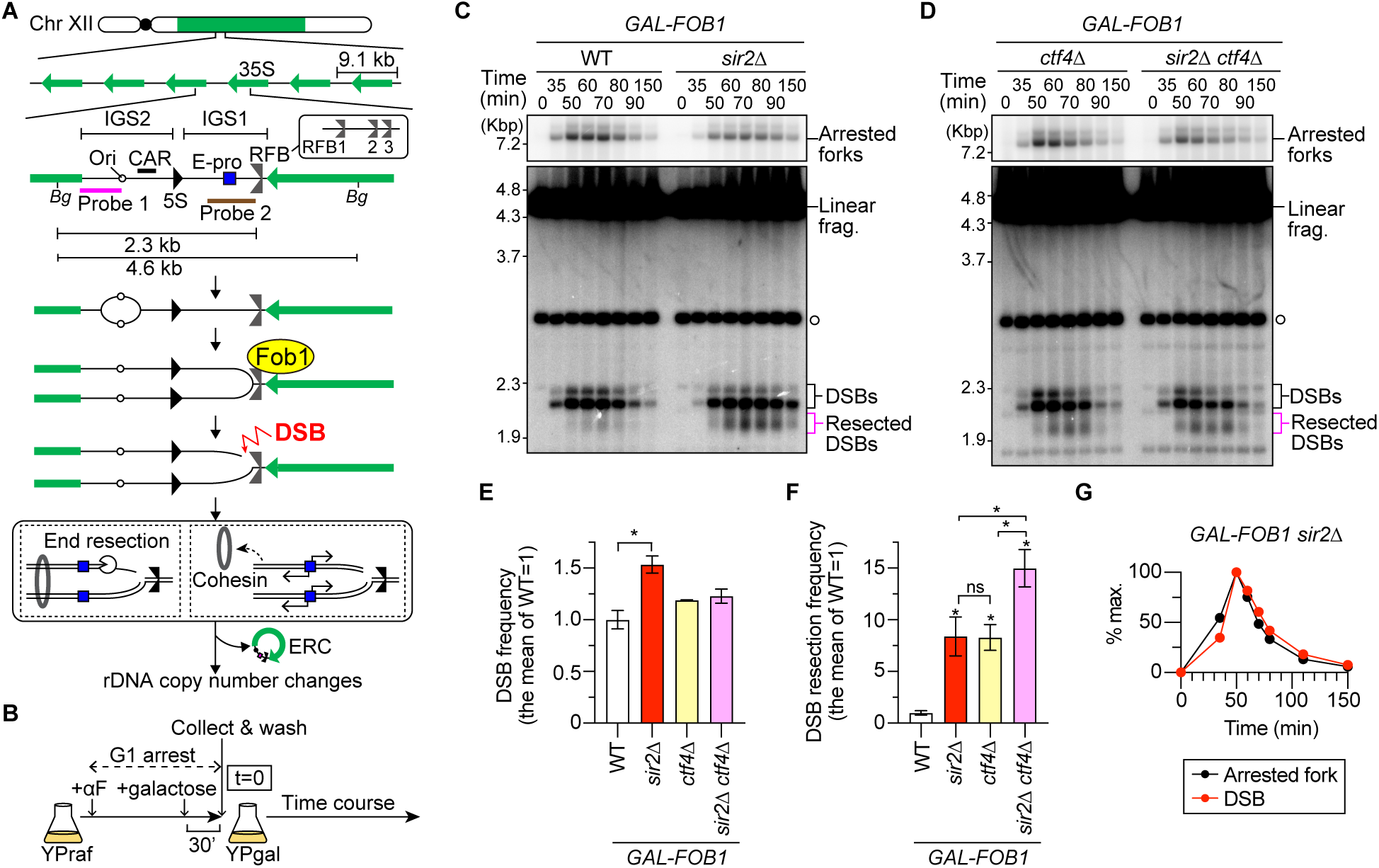
Sir2 suppresses end resection of replication-associated DSBs at the rDNA RFB. **(A)** Replication fork arrest, DSB formation and rDNA copy number changes in the budding yeast rDNA cluster. Chr XII carries the rDNA cluster, as indicated by the green bar. Each copy contains 35S rRNA (35S), 5S rRNA (5S), an origin of DNA replication (ori), a cohesin-associated region (CAR), a bidirectional promoter for noncoding RNA, called E-pro, and a replication fork barrier (RFB). *Bg* indicates a recognition site by the restriction enzyme Bgl II. The pink and brown bars indicate the positions of probe 1 and probe 2 used in the DSB and ERC assays, respectively. (B) Outline of time course experiments to perform DSB assays. (C, D) DSB assay. Cells were collected at different time points in time course experiments after being released from G1 arrest, as described in (B). Genomic DNA was digested with Bgl II, separated by agarose gel electrophoresis, and analyzed by Southern blotting with rDNA probe 1, as indicated in (A). The open circle indicates the terminal fragment that contains the telomere-proximal rDNA repeat and its adjacent non-rDNA fragment. Bands corresponding to arrested forks, linear fragments, DSBs, and resected DSBs are indicated. (E) The frequency of DSBs is expressed as the proportion of DSBs relative to arrested forks and was normalized to the average frequency in the WT. Bars are the mean ± s.e.m. One-way ANOVA was used for multiple comparisons. An asterisk indicates a statistically significant difference (*P* < 0.05) between WT and *sir2Δ*. (F) Frequency of DSB end resection, expressed as the proportion of resected DSBs relative to DSBs and normalized to the average frequency in the WT. Bars are mean ± s.e.m. One-way ANOVA was used for multiple comparisons. Asterisks above the bars and brackets indicate statistically significant differences (*P* < 0.05) between the WT strain and the indicated strains and between different strains, respectively. ns indicates no statistically significant difference. (G) Arrested forks and DSBs are expressed as percentages of the maximum level of DSBs in one representative time course experiment of the *GAL-FOB1 sir2Δ* strain.

Extrachromosomal rDNA circles (ERCs) are produced from the rDNA array (Fig. 1A) (Sinclair and Guarente, 1997), and their level is inversely correlated to the rDNA array size (Mansisidor et al., 2018). Therefore, budding yeast cells have evolved mechanisms to either induce or restrict rDNA copy number changes, depending on the cellular needs, i.e., the number of rDNA copies carried by the cells [reviewed by (Sasaki and Kobayashi, 2023)].

DNA replication forks pause at various roadblocks on template DNA, such as DNA damage, tight protein‒DNA complexes, and transcription machinery [reviewed by (Mirkin and Mirkin, 2007; Branzei and Foiani, 2010)]. Failures to properly resolve stalled forks are a major cause of genome instability. The budding yeast rDNA locus is unique in that once DNA replication is initiated, arrest of the replication fork in the direction opposite to the 35S rDNA is programmed by an rDNA-specific Fob1 protein bound to the RFB site (Fig. 1A) (Brewer and Fangman, 1988; Brewer et al., 1992; Kobayashi et al., 1992). Cleavage of the template of the leading strand at an arrested fork via an as-yet unidentified enzyme leads to the generation of a one-ended DSB (Weitao et al., 2003; Burkhalter and Sogo, 2004; Kobayashi et al., 2004).

Homologous recombination (HR) is often considered an error-free pathway for repairing two-ended DSBs [reviewed by (Paques and Haber, 1999; Symington et al., 2014)]. HR is initiated by degrading the 5’-ended strand of the DSB, the process referred to as DSB end resection. The Mre11-Rad50-Xrs2 (MRX) (a human homolog of Xrs2 is Nbs1) complex is central to HR-mediated repair of two-ended DSBs by initiating DSB end resection [reviewed in (Oh and Symington, 2018; Casari et al., 2019)]. Resected DSBs are subsequently repaired using a homologous sequence on the sister chromatid as a template for Rad52-dependent recombination reactions, which properly restore the original sequence at the DSB site. When DSBs are formed in repetitive regions, however, HR can use a homologous template located at a misaligned position on the sister chromatid or elsewhere in the genome, resulting in chromosome rearrangements [reviewed by (Sasaki *et al*., 2010)].

We previously demonstrated that DSB end resection is normally suppressed for one-ended DSBs formed at Fob1-dependent arrested replication forks at the rDNA RFB, and repair of these DSBs can be completed without Rad52, an essential factor in HR (Sasaki and Kobayashi, 2017). The replisome component Ctf4 is needed for this suppression, and its absence induces DSB end resection, HR-dependent DSB repair and pathological rDNA hyperamplification (Sasaki and Kobayashi, 2017). Interestingly, the suppression of DSB end resection is also relieved when physiological rDNA amplification occurs in the rDNA low-copy-number strain. Thus, the cellular decision to initiate or suppress DSB end resection is the first important regulatory step that impacts the outcome of DSB repair (Fig. 1A) [reviewed by (Sasaki and Kobayashi, 2023)].

Cohesin complexes are recruited to DNA damage sites (Kim et al., 2002; Strom et al., 2004; Unal et al., 2004). Cohesins are associated with the CAR in rDNA and function to suppress rDNA copy number changes by maintaining sister chromatids in close proximity and promoting the use of aligned rDNA copies for DSB repair (Fig. 1A) (Kobayashi *et al*., 2004). A previous study constructed a strain that only carries two rDNA copies, in which E-pro is replaced by the galactose-regulatable, bidirectional *GAL1/10* promoter; this strain is hereafter referred to as the Gal-pro strain (Kobayashi and Ganley, 2005). Transcription from this promoter inhibits cohesin binding to the CAR and induces rDNA amplification, while the opposite is observed when transcription is turned off. The histone deacetylase Sir2, a homolog of mammalian sirtuins, suppresses rDNA copy number changes and extends the replicative lifespan through repression of transcription at the E-pro (Kaeberlein et al., 1999; Kobayashi *et al*., 2004). When the rDNA copy number is reduced, *SIR2* expression decreases, which leads to derepression of transcription at the E-pro, inhibition of cohesin binding to the CAR and induction of rDNA amplification (Kobayashi and Ganley, 2005; Iida and Kobayashi, 2019). Therefore, the regulation of E-pro activity is another process that impacts the fate of DSB repair by influencing cohesin associations with CARs (Fig. 1A).

Here, we showed that in addition to its previously known function in promoting cohesin association, Sir2 is important for suppressing DSB end resection at the RFB. In its absence, DSB end resection initiates HR, which mediates chromosomal rDNA copy number changes and ERC production. The MRX complex was needed for DSB repair. Rad52 was responsible for inducing rDNA copy number changes. Although the absence of Rad52 leads to failure to repair meiotic DSBs and DSBs formed in G2/M, its absence did not compromise the final step of repair of replication-associated DSBs at the rDNA RFB. By turning nearby transcription on or off in the Gal-pro strain, we demonstrated that transcriptional activity near arrested DNA replication forks was responsible for enhancing DSB formation, DSB end resection and HR-mediated repair. Transcription that proceeded toward the RFB was terminated by arrested replication forks, indicating that the collision of transcription machinery with arrested replication forks impacts DSB formation and repair. In addition to being recruited to a previously known site for cohesion recruitment at the CAR, cohesins were recruited to the RFB in an S-phase-specific and Fob1-dependent manner, and cohesin recruitment to both CAR and RFB sites was inhibited by nearby transcription. Taken together, these findings demonstrate that transcription has a substantial impact on the fate of nearby arrested replication forks by inducing DSB formation, DSB end resection and rDNA copy number changes.

## RESULTS

### Sir2 suppresses end resection of replication-associated DSBs in the rDNA

Suppression of DSB end resection is relieved in rDNA low-copy-number strains (Sasaki and Kobayashi, 2017), in which *SIR2* expression is decreased (Iida and Kobayashi, 2019). These findings led us to hypothesize that Sir2 is involved in the suppression of DSB end resection. To test this possibility, we performed time course experiments during synchronous S phase progression in WT and *sir2Δ* strains expressing *FOB1* under the control of a galactose-regulatable *GAL1* promoter (Fig. 1B), followed by the previously developed DSB assay, which detects arrested forks, DSBs and resected DSBs generated at the RFB by Southern blotting (Fig. 1C) (Sasaki and Kobayashi, 2017). Similar to the WT strain, the *sir2Δ* mutant exhibited S-phase-specific replication fork arrest and DSB formation at the RFB sites. At the RFB, there are three sites, RFB1–3, that can arrest the progression of replication forks (Fig. 1A) (Brewer *et al*., 1992; Kobayashi, 2003): RFB1 exhibits stronger activity in arresting replication forks than the other two sites that are very close to each other. Thus, the lower DSB band corresponds to DSBs formed at RFB1, while the upper DSB band corresponds to DSBs formed at RFB2 and 3 (Fig. 1C). Deletion of *SIR2* led to an ∼1.5-fold increase in the frequency of DSBs compared to that in the WT strain (Fig. 1C, 1E). Such an increase was not observed in a previous study (Kobayashi *et al*., 2004), most likely because asynchronous cultures were used and the peak of DSB formation was missed. Thus, Sir2 functions to reduce DSB formation at arrested forks.

The absence of Sir2 led to an increase in the frequency of DSB end resection by ∼8.4-fold compared to that in the WT cells, which was comparable to the frequency observed in cells lacking Ctf4 (Fig. 1C, 1D, 1F). The frequency of DSB end resection was further greater in the *sir2Δ ctf4Δ* double mutant than in the single mutants, indicating that two different pathways induce DSB end resection (Fig. 1F). Notably, the signal intensities of the resected DSBs in the same membrane were comparable between the *ctf4Δ* and *sir2Δ ctf4Δ* mutants (Fig. 1D).

However, when we normalized the level of resected DSBs to the level of DSBs, the frequency of DSB end resection was reproducibly greater for the *sir2Δ ctf4Δ* mutant than for the *sir2Δ* and *ctf4Δ* single mutants (Fig. 1F). Therefore, Sir2 suppresses end resection of DSBs formed at the RFB, but its action is distinct from that mediated by Ctf4.

### The MRX complex but not Rad52 is required for DSB repair in the *sir2Δ* mutant

Deletion of *MRE11* from the *sir2Δ* strain suppressed end resection of one-ended DSBs formed at the RFB by >100-fold compared with that in the *sir2Δ* single mutant (Fig. 2A, 2C). Therefore, the MRX complex is responsible for initiating DSB end resection in the *sir2Δ* mutant. Compared with the *sir2Δ* mutant, the *sir2Δ mre11Δ* strain accumulated DSBs at the end of the time course, suggesting that DSB repair was severely compromised (Fig. 1C, 1G, 2A, 2B). The *sir2Δ mre11Δ* strain exhibited severe growth defects, which were substantially suppressed by inactivation of Fob1 (Fig. 2D, *sir2Δ mre11Δ FOB1* vs. *sir2Δ mre11Δ fob1*), consistent with a previous finding (Shyian et al., 2016). Therefore, the MRX complex is crucial for DSB repair at the RFB, the deficiency of which compromises cellular growth.

**Figure 2.**
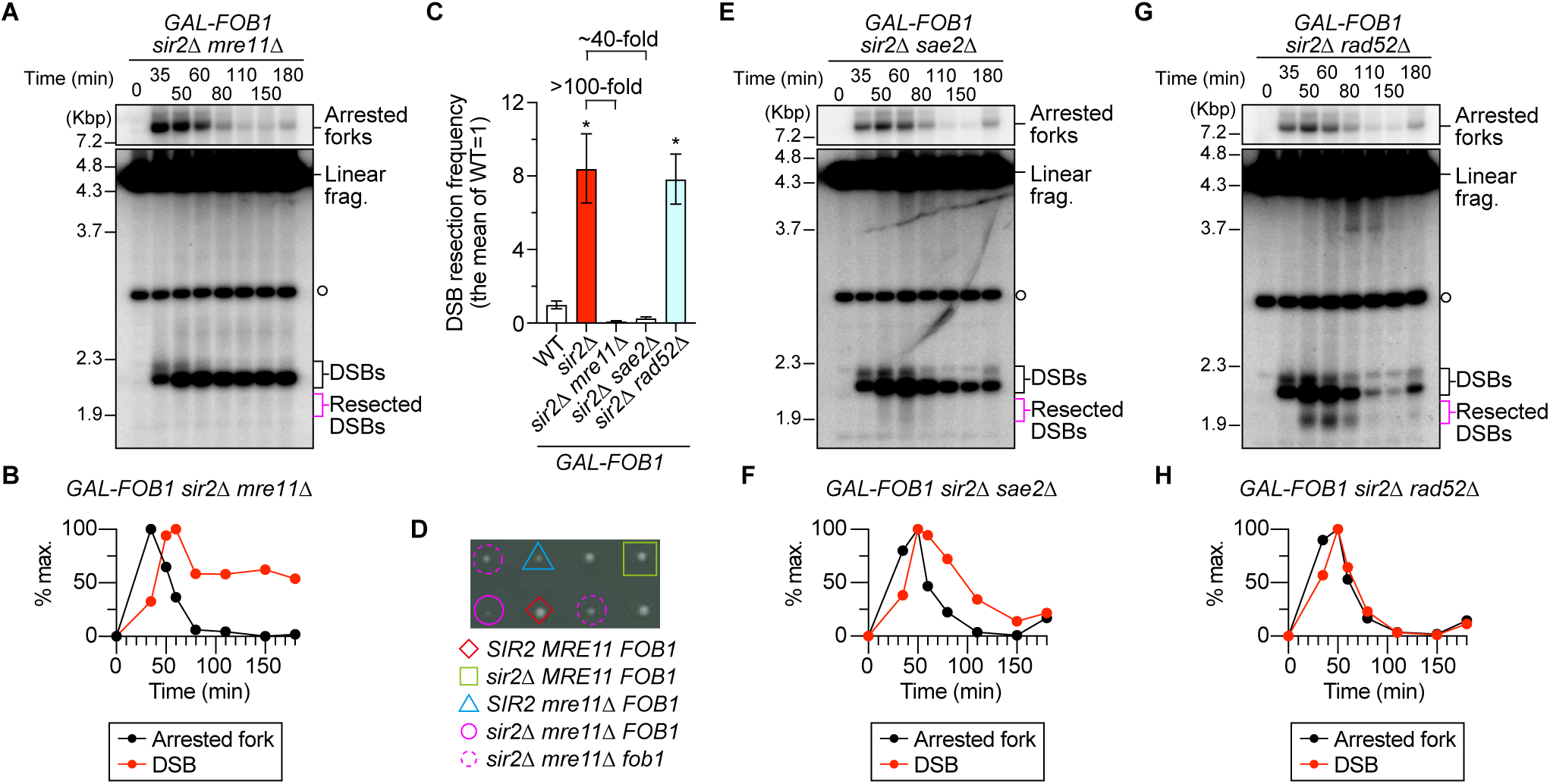
The MRX complex but not Rad52 is needed for DSB repair at the RFB in the *sir2Δ* mutant. (A, E, G) DSB assay. The time course and DSB assays were performed as described for Fig. 1B and 1C. The terminal fragment, arrested forks, linear fragments, DSBs, and resected DSBs are indicated. (B, F, H) Arrested forks and DSBs are expressed as percentages of the maximum values in one representative time course experiment of the indicated strains. (C) Frequency of DSB end resection, determined as described for Fig. 1F. The bars are the means ± s.e.m. One-way ANOVA was used for multiple comparisons. Asterisks above the bars indicate statistically significant differences (*P* < 0.05) between the WT and the indicated strains. (D) Colony sizes of the indicated strains. Haploid clones of different genotypes were isolated by tetrad dissection of diploids heterozygous for *sir2Δ*, *mre11Δ*, and *fob1*.

Sae2 (CtIP in humans) is known as a cofactor for Mre11’s catalytic activity (Cannavo and Cejka, 2014; Cannavo et al., 2019). Deletion of *SAE2* from the *sir2Δ* strain caused an ∼40-fold reduction in the number of resected DSBs but did not impair DSB repair (Fig. 2C, 2E, 2F). Furthermore, the DSB signal disappeared at the normal time point in the *sir2Δ* strain lacking Rad52 (Fig. 2G, 2H). These findings demonstrate that the MRX complex is essential for DSB end resection and repair but that resected DSBs can be repaired without the assistance of Rad52 in cells lacking Sir2. Because the *ctf4Δ* strain requires Rad52 for DSB repair (Sasaki and Kobayashi, 2017), the manner in which DSBs are repaired in the *sir2Δ* strain is distinct from that in the *ctf4Δ* strain.

### Rad52 mediates rDNA copy number changes during DSB repair

Rad52 is responsible for mediating rDNA amplification in a low-copy rDNA strain in which the *SIR2* expression level is decreased (Kobayashi *et al*., 2004; Iida and Kobayashi, 2019). This finding was interpreted to indicate that HR is essential for the repair of DSBs formed at the RFB. However, this study demonstrated that Rad52 was not essential for DSB repair in the *sir2Δ* mutant (Fig. 2G, 2H). To address these confounding observations, we tested whether HR is indeed involved in rDNA copy number changes in the *sir2Δ* mutant.

The frequency of chromosomal rDNA copy number changes can be assessed by analyzing the shape of the band of chr XII by pulsed-field gel electrophoresis (PFGE). The *sir2Δ* mutant undergoes expansion and contraction of the rDNA array, causing a smeared chr XII band (Fig. 3A), consistent with previous finding (Kobayashi *et al*., 2004; Goto et al., 2021; Yokoyama et al., 2023). The viability of cells lacking both Sir2 and Mre11 was compromised due to DSB repair defects at the RFB, but in the surviving cells, DSB end resection was defective (Fig. 2A– 2D). In these surviving cells, the chr XII band became sharper, indicating that chromosomal rDNA copy number changes were suppressed (Fig. 3A). ERCs are produced from the rDNA region, but its formation was also suppressed in the *sir2Δ mre11Δ* mutant (Fig. 3B, 3C). Moreover, deletion of *RAD52* suppressed both chromosomal rDNA copy number changes and ERC production in the *sir2Δ* strain (Fig. 3D–3F). Therefore, Rad52 is responsible for inducing rDNA copy number changes during DSB repair in the *sir2Δ* mutant, but cells have a backup system to complete DSB repair without HR (see discussion).

**Figure 3.**
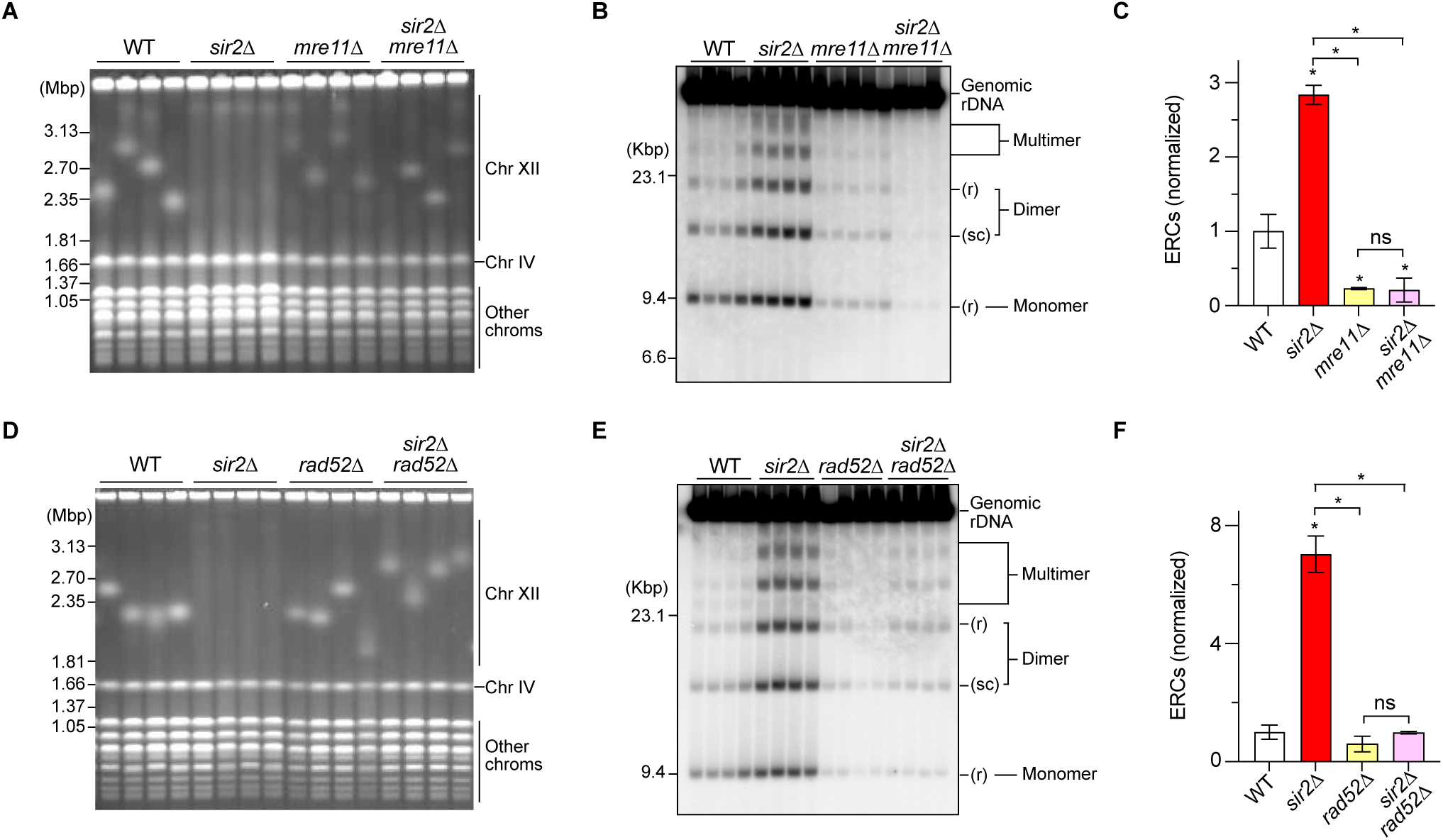
The MRX complex and Rad52 are responsible for rDNA copy number changes in the *sir2Δ* mutant. (A, D) PFGE analysis to examine the size heterogeneity of chr XII. DNA from independent clones of the indicated strains was separated by PFGE and stained with ethidium bromide. M indicates *Hansenula wingei* chromosomal DNA markers. (B, E) ERC detection. DNA was separated by agarose gel electrophoresis, followed by Southern blotting with rDNA probe 2, as shown in Fig. 1A. Genomic rDNA and different forms of ERCs are indicated. r and sc indicate relaxed and supercoiled forms of ERCs, respectively. M indicates λ DNA-Hind III markers. (C, F) The level of ERCs. The sum of monomers, dimers, and multimers was determined relative to genomic rDNA and normalized to the average of the WT clones (bars show the mean ± s.e.m.). One-way ANOVA was used for multiple comparisons. Asterisks above the bars and brackets indicate statistically significant differences (*P* < 0.05) between the WT strain and the indicated strains and between different strains, respectively. ns indicates no statistically significant difference.

### Sir2 mediates cohesin recruitment to arrested replication forks

To elucidate how Sir2 suppresses rDNA copy number changes, we revisited the previous finding that Sir2 promotes cohesin associations with CAR in exponentially growing cells (Laloraya et al., 2000; Kobayashi *et al*., 2004; Kobayashi and Ganley, 2005). We examined how Sir2-mediated regulation of transcription from E-pro regulates cohesin associations during S phase when DSB repair occurs and how the expression of Fob1, which is responsible for programmed replication fork arrest, influences cohesin binding to rDNA. To this end, we performed time course experiments by arresting cells in G1 and subsequently releasing the cells into media supplemented with nocodazole to arrest the cells in M phase and performing chromatin immunoprecipitation of a subunit of a cohesin complex, Mcd1, in the strain in which *MCD1* was tagged with a Flag tag in the *GAL-FOB1* background (Fig. 4A).

**Figure 4.**
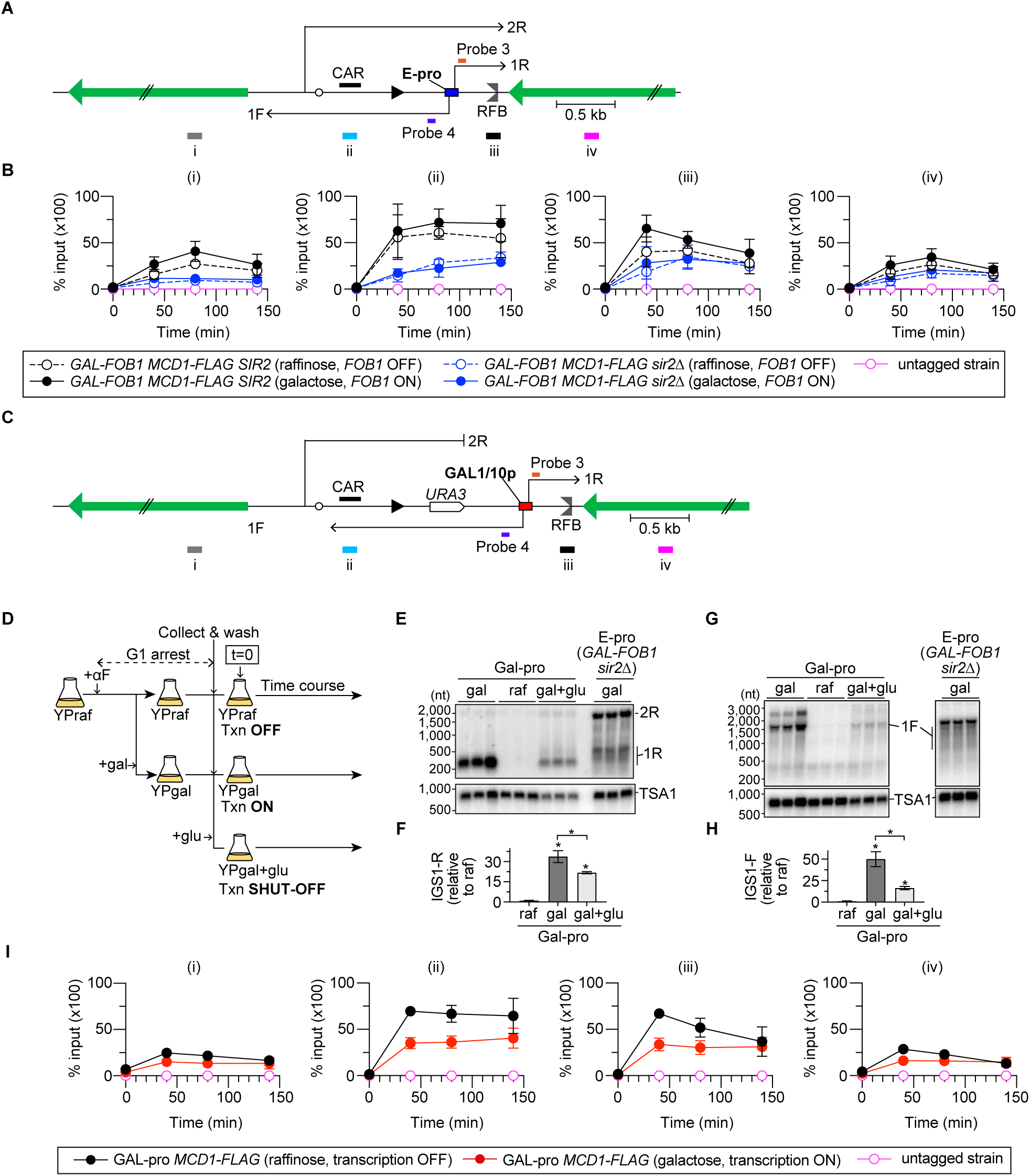
Cohesin associations with rDNA are differentially regulated by transcription and replication fork arrest. (A, C) The regions in the rDNA where cohesin enrichments were examined in the E-pro (A) and Gal-pro (C) strains. Probes 3 and 4 are single-stranded DNA probes used to detect noncoding RNA transcribed from the E-pro (A) and Gal-pro (C) strains. Four different regions (i)–(iv) were examined. (B, I) ChIP‒qPCR analyses of Mcd1-Flag in the indicated strains grown in the presence of the indicated carbon sources. Untagged strains were used as a control. Time courses were performed as indicated in Fig. 1B and Fig. 4D, except that cells were released from G1 arrest into the nocodazole-containing media to arrest cells in the G2/M phase of the cell cycle. ChIP was performed using an antibody against a Flag epitope tag. The enrichment of chromatin in the indicated regions is expressed as the % input (IP/whole-cell extract). The bars show the means ± s.e.m. from three independent cultures of each strain, except for the untagged strain, which indicates the range from two independent experiments. (D) Outline of time course experiments for ChIP of the *Gal-pro MCD1-FLAG* strain and Northern blotting and DSB assays of the *Gal-pro* strains. (E, G) Detection of noncoding RNAs. IGS1-R (1R) and IGS2-R (2R) (E) and IGS1-F (1F) (G) transcribed from the Gal-pro strain grown under different conditions and from the *GAL-FOB1 sir2Δ* strain were detected by Northern blotting. RNA samples isolated at t=40 time points were analyzed. The membranes were reprobed for *TSA1* transcripts as a loading control. (F, H) Levels of IGS1-R (F) and IGS1-F (H). The level of transcripts was quantified relative to that of *TSA1*, which was normalized to the average level of transcripts in the Gal-pro strain grown in the presence of raffinose. The bars show the means ± s.e.m. One-way ANOVA was used for multiple comparisons.

In the absence of *FOB1* expression in *SIR2* cells, cohesin was most enriched at the CAR in the rDNA during S phase and remained associated until G2/M (Fig. 4A, 4B, ii, open black circle). The expression of *FOB1* did not substantially alter cohesin association with the CAR (Fig. 4A, 4B, ii, closed black circle). In contrast to the CAR, *FOB1* expression led to a prominent increase in the association of cohesin with RFB during S phase, and this association decreased as the cells progressed toward the G2/M phase (Fig. 4B, iii, open vs. closed black circles). The S-phase specific, Fob1-dependent binding of cohesin to RFB was not observed in previous studies (Laloraya *et al*., 2000; Kobayashi *et al*., 2004; Kobayashi and Ganley, 2005), most likely because those studies analyzed cohesin associations from exponentially growing cultures where cells were mostly in G1 or G2/M phase. Our findings demonstrated that in addition to the previously known cohesin binding at the CAR, cohesin is recruited to the RFB site during S phase in response to *FOB1* expression.

Cohesin association was inhibited at the CAR when Sir2 was absent, but this association was not altered by *FOB1* expression (Fig. 4B, ii). This result extended previous findings and demonstrated that Sir2 promotes cohesin binding to the CAR (Kobayashi *et al*., 2004), regardless of the presence or absence of Fob1. Importantly, Fob1-dependent cohesin association at the RFB was not observed when Sir2 was absent (Fig. 4B, iii, open vs. closed blue circles). Therefore, Sir2 mediates cohesin recruitment to RFB in response to Fob1-dependent replication fork arrest and/or DSB formation.

To test whether the transcriptional activity of E-pro regulates cohesin binding to rDNA, cohesin associations were analyzed in the Gal-pro strain, in which E-pro was replaced by the *GAL1/10* promoter in the rDNA two-copy strain and rDNA copies were amplified to ∼80 copies (Kobayashi and Ganley, 2005). IGS1-R transcripts that were synthesized toward RFB were barely detected in the presence of raffinose but were elevated by the addition of galactose to the media (Fig. 4E, 4F). These transcripts were relatively short compared with the IGS1-R transcripts synthesized from the native E-pro in the *GAL-FOB1 sir2Δ* strain (Fig. 4E). IGS1-F transcripts that were synthesized toward 5S rDNA were also induced by the addition of galactose to the media, although these transcripts were longer due to the presence of the *URA3* gene (Fig. 4C, 4G, 4H). Thus, the transcription activity near the RFB can be turned on or off in the Gal-pro strain by the addition of galactose or raffinose, respectively, to the media.

When transcription was kept OFF, the Gal-pro cells exhibited cohesin binding to the CAR during S phase, which lasted until G2/M phase (Fig. 4C, 4I, ii, black circle). In these cells, cohesin was also recruited to the RFB during S phase but decreased as the cells progressed toward G2/M (Fig. 4I, iii, black circle). These cohesin association patterns were similar to those observed in *GAL-FOB1 SIR2* cells expressing *FOB1* (Fig. 4B, 4I). In contrast, induction of transcription from the *GAL1/10* promoter substantially decreased the level of cohesin bound not only to the CAR but also to the RFB site, mimicking the binding patterns observed in the *GAL-FOB1 SIR2* vs. *sir2Δ* strains expressing *FOB1* (Fig. 4B, 4I). Therefore, Sir2-mediated repression of transcription from E-pro enables cohesin binding to CAR and RFB in a manner independent and dependent on Fob1, respectively. Considering that a temperature-sensitive *smc1* mutant deficient in another cohesin subunit shows an increased frequency of loss of a marker gene from the rDNA cluster (Kobayashi et al., 2004), cohesins bound to the CAR and RFB both contribute to restricting the interaction of the DSB end with the misaligned rDNA copy during DSB repair.

### Transcription enhances DSB formation and end resection at RFBs

To examine how the transcription status of E-pro affects the fate of arrested replication forks, we monitored DSB formation and repair in the Gal-pro strain (Fig. 4D, 5A). The Gal-pro cells displayed two classes of Fob1-and S-phase-specific DSB band signals (Fig. 5B and 5E; Gal-pro *FOB1* vs. *fob1*), among which the lower band, which was homogeneous in size, most likely corresponded to DSBs formed at RFB1, and the upper smeared signal was derived from DSBs formed at RFB2 and RFB3 (Fig. 1A, 5B, 5E). Transcription activity altered the pattern of DSB formation by rendering more frequent DSB formation at the RFB1 site than at the RFB2 and RFB3 sites (Fig. 5B, 5E, raf vs. gal in the Gal-pro *FOB1* strain). Transcription induction not only elevated the DSB frequency by 40% (Fig. 5C) but also caused an increase in the frequency of DSB end resection (Fig. 5D). Therefore, transcription near the RFB stimulates DSB formation as well as its end resection.

**Figure 5.**
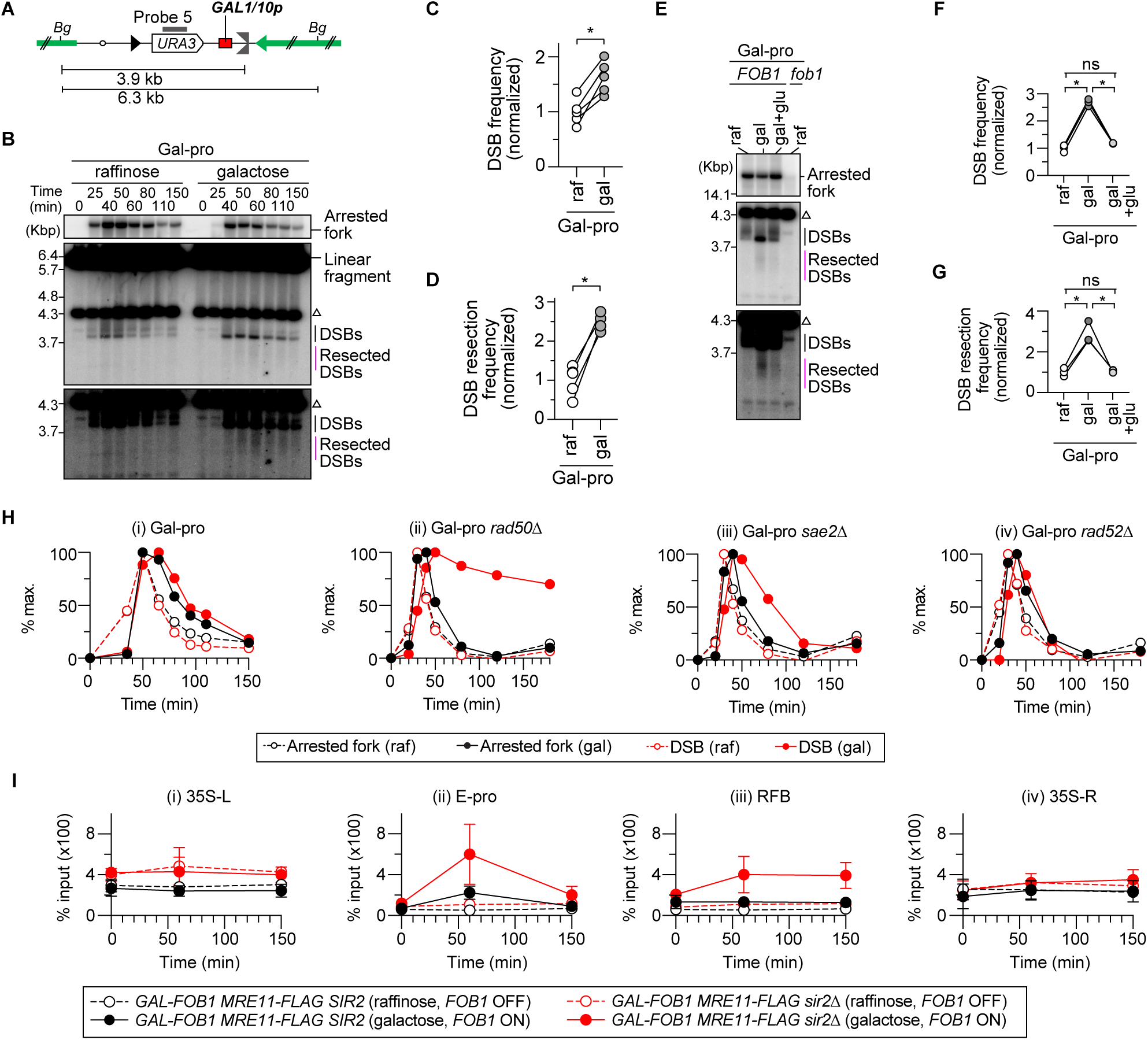
Transcription activity near the RFB site induces MRX-dependent DSB repair. (A) The position of probe 5 used for DSB assays with the Gal-pro strains. *Bg* indicates a recognition site by the restriction enzyme Bgl II. (B, E) DSB assays. The time course and DSB assays were performed as described for Fig. 1B and 1C. Arrested forks, linear fragments, DSBs, and resected DSBs are indicated. The triangle indicates the Bgl II fragment around the endogenous *URA3*. In (E), arrested forks, DSBs, and resected DSBs were analyzed in samples of the Gal-pro strain and Gal-pro strain lacking Fob1 grown in the presence of the indicated carbon sources. DNA samples at the time points where the DSB level reached a maximum in (B) were analyzed side-by-side. The lower panels show a more exposed contrast image of the phosphor imager signal around DSBs. (C, F) The frequency of DSBs is expressed as the proportion of DSBs relative to the number of arrested fork signals and was normalized to the average frequency of DSBs in the Gal-pro strain grown in raffinose-containing media. The bars are the means ± s.e.m. Two-sided Student’s t tests were used to compare DSB frequencies between cells grown in the presence of raffinose and galactose. An asterisk indicates a statistically significant difference (*P* < 0.05). (D, G) Frequency of DSB end resection, expressed as the proportion of resected DSBs relative to DSBs, which was normalized to the average level of DSB frequency in the Gal-pro strain grown in raffinose-containing media. The bars are the means ± s.e.m. Two-sided Student’s t tests were used to compare DSB frequencies between cells grown in the presence of raffinose and galactose. An asterisk indicates a statistically significant difference (*P* < 0.05), and ns indicates no statistically significant difference. (H) Arrested forks and DSBs are expressed as percentages of the maximum values in one representative time course experiment of the indicated strains. (I) ChIP‒qPCR analyses of Mre11-Flag in the indicated strains grown in the presence of the indicated carbon sources. The time course was performed as indicated in Fig. 1B. ChIP was performed using an antibody against a Flag epitope tag. The enrichment of chromatin in the indicated regions is expressed as the % input (IP/whole-cell extract). The bars show the mean ± s.e.m. from three independent cultures of each strain.

To test which aspects of transcription influence DSB formation and its end resection, we monitored these processes in the condition where *de novo* transcription was shut off at the beginning of time courses by the addition of glucose, which causes rapid transcriptional repression from the *GAL1/10* promoter (Fig. 4D). The levels of transcripts in cells where transcription was inhibited were reduced by ∼1.6–3.3-fold compared to those in cells grown in galactose-containing media (Fig. 4E–4H, gal vs. gal+glu), while the levels were still ∼10–20-fold greater than those in cells grown in raffinose-containing media (Fig. 4E–4H, raf vs. gal+glu). Inhibition of *de novo* transcription led to suppression of DSB formation and its end resection almost down to their frequencies seen in the raffinose-containing media (Fig. 5E–5G). Thus, it is less likely that the IGS1-R and IGS1-F transcripts that are released from the template impact DSB formation and end resection. Instead, these processes are affected by the presence of transcription machinery on DNA or other transcription-associated events, such as R-loops, which are formed during transcription and consist of RNA‒DNA hybrids and a displaced DNA strand (see discussion) (Garcia-Muse and Aguilera, 2019; Petermann et al., 2022).

### The MRX complex is essential for transcription-induced DSB repair at RFBs

We next examined how transcriptional activity influences DSB repair. In the Gal-pro strain, the timing of DSB signal disappearance was comparable, regardless of whether transcription at the *GAL1/10* promoter was turned on or off (Fig. 5H, i). In the Gal-pro strain lacking Rad50, DSBs were promptly repaired when transcription was kept off. In striking contrast, when transcription was induced, the Gal-pro *rad50Δ* strain exhibited severe delays in DSB repair (Fig. 5H, ii). Therefore, the transcriptional activity near the RFB renders DSB repair dependent on the MRX complex.

When transcription was activated near the RFB, the Gal-pro *sae2Δ* strain exhibited a slight delay in the disappearance of DSB signals but was proficient at repairing DSBs by the end of the time course (Fig. 5H, iii). Finally, DSB repair occurred at the normal timing in the Gal-pro strain lacking Rad52, regardless of the transcriptional activity near the RFB (Fig. 5H, iv). The genetic requirements for DSB repair in the transcriptionally active Gal-pro strain were similar to those observed in the *GAL-FOB1 sir2Δ* background (Fig. 2A, 2B, 2E–2H, 5H), in which transcription from E-pro was derepressed.

The MRX complex did not show strong enrichment across the rDNA sequence when programmed replication fork arrest was inhibited by the *GAL-FOB1* strain not expressing *FOB1*, regardless of the presence or absence of Sir2 (Fig. 5I, *FOB1* OFF). When *FOB1* was expressed to induce replication fork arrest in *SIR2* cells, we did not observe substantial binding of the MRX complex to RFB (Fig. 5I, iii). However, the MRX complex was recruited to RFB in *sir2Δ* cells during S phase, which remained associated until the end of the time course experiments, when most cells were in G2/M (Fig. 5I, iii).

In *SIR2* cells, the MRX complex was associated with E-pro during S phase upon *FOB1* expression, and this association decreased toward G2/M (Fig. 5I, ii). The level of the MRX complex bound to E-pro was substantially increased in *sir2Δ* cells that expressed *FOB1* (Fig. 5I, *GAL-FOB1 sir2Δ*, *FOB1* ON). Therefore, the co-occurrence of replication fork arrest and transcription from E-pro recruit the MRX complex to E-pro and less to RFB, consistent with the dependency of the MRX complex on DSB repair in the transcriptionally active region (Fig. 5H, 5I).

### Transcription is terminated by arrested forks at the RFB

We tested whether replication fork arrest and/or DSB formation impact transcription from E-pro (Fig. 6A). The expression of *FOB1* in *SIR2* cells caused an ∼2-fold reduction in the IGS1-F transcript level throughout the cell cycle compared to that in cells not expressing *FOB1* (Fig. 6B, 6C). A reduction in the level of IGS1-F transcripts was also observed upon *FOB1* expression in *sir2Δ* cells (Fig. 6D, 6E). Considering that transcription from the E-pro locus inhibits cohesin association with CAR (Fig. 4B, 4I) (Kobayashi and Ganley, 2005), why did Fob1-dependent reduction in IGS1-F transcripts not lead to elevation of cohesin association with CAR upon *FOB1* expression in *SIR2* and *sir2Δ* cells (Fig. 4B, 6B–6E)? IGS1-F transcripts can be degraded by the Ccr4/Pop2/Not mRNA deadenylase complex (Hosoyamada et al., 2019). Therefore, we speculate that the ability of IGS1-F to be transcribed is comparable in cells with or without *FOB1* expression, that IGS1-F transcripts are degraded more efficiently in the presence of Fob1, and that cohesin associations are not affected by the presence or absence of *FOB1* expression.

**Figure 6.**
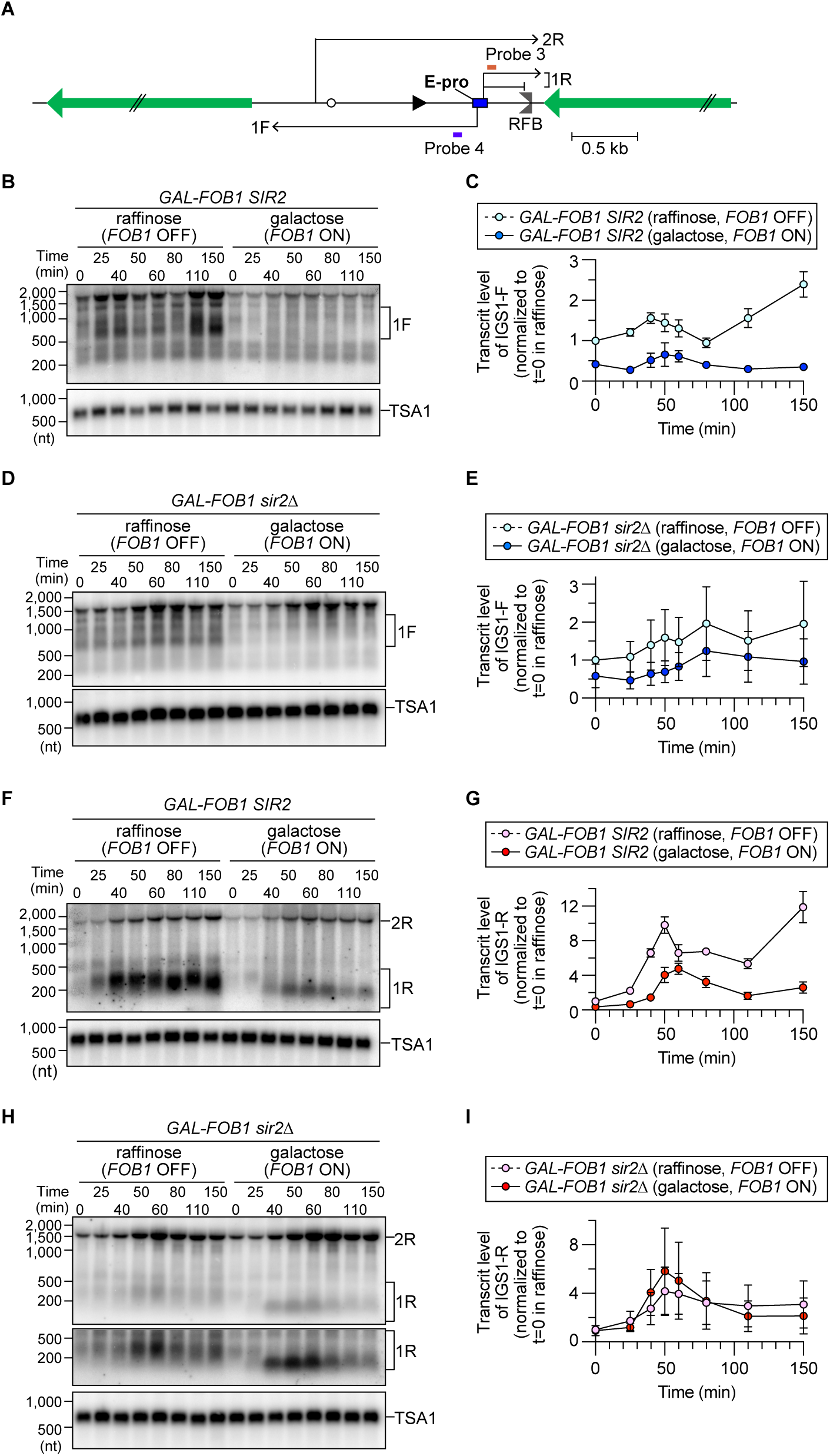
Fob1 protein causes premature termination of transcription from E-pro. (A) The positions of the probes and direction of transcription from E-pro. Probes 3 and 4 are single-stranded DNA probes used to detect noncoding RNA transcribed from E-pro. (B, D, F, H) Detection of IGS1-F (1F) (B, D), IGS1-R (1R) and IGS2-R (2R) (F, H). Time courses were performed for the indicated strains grown in the presence of raffinose or galactose, as described in Fig. 1B. Total RNA was isolated and analyzed by Northern blotting with single-stranded DNA probes, as indicated in Fig. 5A. The membranes were reprobed for *TSA1* transcripts as a loading control. (C, E, G, I) Levels of IGS1-F (B, D) and IGS1-R (F, H). In each experiment, the level of the transcript was determined relative to that of *TSA1* at each time point. The fold increase in the transcript level was subsequently determined relative to that at t=0 in the raffinose-containing culture. The bars show the range from two independent experiments.

*FOB1* expression caused a reduction in the level of IGS1-R transcripts in *SIR2* cells (Fig. 6F, 6G). However, the level of IGS1-R transcripts was not altered by *FOB1* expression in *sir2Δ* cells (Fig. 6H, 6I). Thus, in contrast to the transcription pattern of IGS1-F, Sir2 is involved in lowering the level of IGS1-R transcripts upon *FOB1* expression.

In *SIR2* cells expressing *FOB1*, the IGS1-R transcripts generated at t=0 and 25 min were similar in length to those synthesized in *SIR2* cells not expressing *FOB1* (Fig. 6F). Interestingly, these transcripts became shorter after 40 min, when the number of arrested replication forks normally peaked (Fig. 6F). This Fob1-mediated shortening of IGS1-R transcripts was also observed in *sir2Δ* cells (Fig. 6H). A previous study demonstrated that the major transcription initiation site for IGS1-R is ∼175 nt away from the RFB (Houseley et al., 2007), indicating that transcription of IGS1-R proceeds beyond the RFB in *FOB1*-expressing cells at early time points but terminates around the RFB when the replication fork is arrested. A previous ChIP study demonstrated that RNAP II is enriched at the RFB (Houseley *et al*., 2007). These findings suggest that the arrest of replication forks caused by Fob1 blocks the progression of RNAP II, which synthesizes IGS1-R.

## DISCUSSION

Our findings demonstrated that transcription near arrested replication forks has a strong impact on both transcription and DNA replication (Figs. 1, 5, 6). The collision of transcription machinery components with arrested replication forks causes termination of transcription (Fig. 6). Furthermore, this collision stimulates not only DSB formation at arrested replication forks but also DSB end resection and subsequent HR-mediated reactions that are prone to chromosomal DNA copy number changes and the production of extrachromosomal circular DNA (Figs. 1, 3, 5, 7A). We offer evidence that the MRX complex becomes essential for DSB repair when DSBs are formed at arrested replication forks in transcriptionally active regions (Figs. 2, 5).

**Figure 7.**
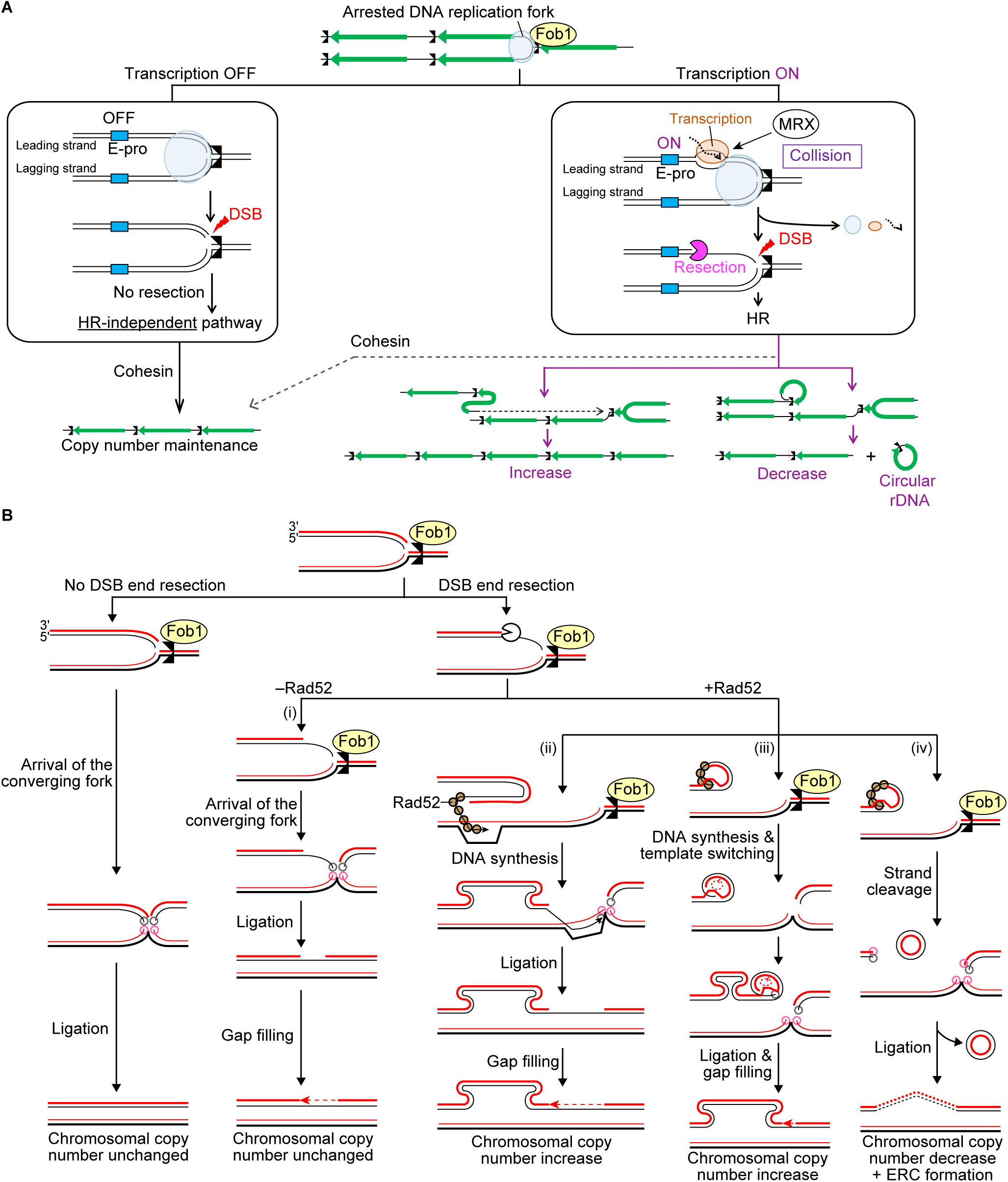
Transcription activity near arrested replication forks is prone to DNA copy number changes and circular DNA formation. (A) A DNA replication fork is arrested at the RFB. (Left) When transcription from E-pro is inactive, arrested forks can be converted into one-ended DSBs, but these DSBs can be repaired by an HR-independent pathway, leading to no change in rDNA copy number. (Right) When transcription from E-pro is activated, the transcription machinery collides with the arrested DNA replication forks. This collision generates R-loop or DNA torsional stress, promoting DSB formation and recruiting the MRX complex. MRX-dependent resolution is accompanied by the initiation of DSB end resection. Cohesin can act post-DSB end resection, ensuring the use of the rDNA copy at an aligned position and the maintenance of rDNA copy number. When cohesin binding is inhibited, DSB repair can be accompanied by chromosomal rDNA copy number changes and ERC formation. (B) Model of the repair of one-ended DSBs formed at arrested replication forks at the RFB. Grey and pink circles indicate two ends of DNA strands that are likely to be joined by ligases. See the text for details.

Budding yeast cells turn on the systems to either stabilize or destabilize the rDNA, depending on the cellular needs [reviewed in (Sasaki and Kobayashi, 2023)]. Our previous study suggested that the stability of the rDNA cluster is dictated by the decision as to whether cells undergo end resection of one-ended DSBs formed at arrested replication forks (Sasaki and Kobayashi, 2017; 2023): when the maintenance of rDNA stability is favored, DSB end resection is suppressed; when the rDNA copy number needs to be changed, the suppression of DSB end resection is relieved. In this study, we reinforce this view and identify Sir2 as a factor involved in suppressing DSB end resection to restrict HR-mediated DSB repair, which leads to chromosomal rDNA copy number changes and ERC production (Figs. 1, 3).

Sir2 has a previously known function in promoting sister chromatid cohesion to CAR (Kobayashi *et al*., 2004). We also revealed that Sir2 is involved in the recruitment of cohesin to RFB in an S-phase specific, Fob1-dependent manner (Fig. 4B). Recruitment of cohesins to both regions may be important for facilitating the use of the rDNA copy at an aligned position on the sister chromatid as a repair template for DSB repair (Fig. 7A). Alternatively, cohesins that are recruited to RFB may have direct roles in facilitating DSB repair (Kim *et al*., 2002; Strom *et al*., 2004; Unal *et al*., 2004).

A deficiency in HR-mediated repair of DSBs that are formed in the G2/M phase in mitotic cells or in meiotic cells compromises cellular viability [reviewed by (Paques and Haber, 1999; Symington *et al*., 2014)]. In striking contrast, the absence of Rad52 in the *sir2Δ* cells did not impair the repair of one-ended DSBs formed at arrested replication forks, although these DSBs were end-resected (Fig. 2G, 2H). In our previous study, we showed that when the replisome factor Ctf4 is absent, DSBs are resected, and repair of these cells requires Rad52 (Sasaki and Kobayashi, 2017). Therefore, the absence of Ctf4 and Sir2 causes DSB end resection and repair in distinct manners. Sir2 regulates these processes by regulating E-pro activity (Figs. 1, 5). On the contrary, we speculate that Ctf4 ensures the stability of the replisome arrested at the RFB.

How are resected DSBs repaired in the absence of Rad52? We propose that in *sir2Δ rad52Δ* cells, resected DSB ends do not undergo strand invasion but instead remain associated with other nonreplicated strands (Fig. 7B, i). When the other replication fork arrives from the downstream origin, nascent strands can be ligated by ligases, for example, DNA ligase I at the replisome, as such ligation normally occurs during Okazaki fragment maturation and during replication termination when two replication forks converge. Then, the resulting gap is filled, and DSB repair is completed. In this case, the rDNA copy number is unchanged (Fig. 7B, i). When Sir2 is absent but Rad52 is present (Fig. 7B, ii–iv), Rad52 binds to the resected DSB end and catalyzes annealing to or invasion into a misaligned rDNA copy. DNA synthesis from the DSB end leads to amplification of the copy but is likely stopped by the Fob1 protein, which remains bound to the RFB site (Fig. 7B, ii). After the converging fork reaches the RFB site, nascent strands are ligated, and a gap is filled (Fig. 7B, ii). In this manner, Rad52 catalyzes rDNA copy number changes but is not needed for final sealing of the broken end. When the DSB end interacts with the other copy on the same chromosome, continuous rounds of DNA synthesis and template switching result in rDNA amplification (Fig. 7B, iii). Alternatively, DNA strands can be clipped to release rDNA copies as an ERC, and subsequent ligation of DNA strands leads to a decrease in chromosomal rDNA copy number (Fig. 7B, iv). Furthermore, Mansisidor et al. (2018) proposed another model for ERC formation in which the formation of DSBs at two RFBs in nearby rDNA copies releases rDNA fragments, and subsequent circularization results in the production of ERCs.

When transcription is low near the arrested replication fork, DSBs are repaired by an HR-independent pathway (Fig. 5H, 7A left and 7B left). We propose that these DSBs are associated with the other nonreplicated strands and ligated to the other double-stranded DNA ends when the converging fork arrives (Fig. 7A left and 7B, left). These models need to be investigated in future studies.

In terms of the relationship between rDNA recombination and transcription, the findings from DSB assays using the Gal-pro strain clearly demonstrated that the MRX complex is needed for DSB repair only when transcription at this locus is high (Fig. 5H). Similarly, the MRX complex is important for DSB repair in the *sir2Δ* mutant, in which transcription from E-pro is activated (Fig. 2A, 2B). Consistent with the importance of the MRX complex for DSB repair, this complex is recruited to E-pro and RFB in the *sir2Δ* mutant in an S-phase-and Fob1-dependent manner (Fig. 5I).

When transcription is activated, the progression of transcription machinery is blocked by replication forks that are physically arrested at the RFB (Fig. 7A, right). Transcription may preferentially occur on a sister chromatid that synthesizes the leading strand because the other sister chromatid that generates the lagging strand may not be a good template for transcription due to the discontinuity of Okazaki fragment. At the rDNA RFB, the template for the leading strand is broken to generate a one-ended DSB (Burkhalter and Sogo, 2004). We speculate that collision of the transcription machinery with the arrested forks on the leading strand is responsible for breakage of the leading strand template. Collision between transcription and arrested replication forks potentially generates R-loops or DNA topological stresses [reviewed in (Bermejo et al., 2012; Aguilera and Gaillard, 2014; Garcia-Muse and Aguilera, 2016; Goehring et al., 2023)]. These stresses stimulate DSB formation (Figs. 1E, 5C). Furthermore, the MRX complex is recruited to resolve these aberrant structures possibly by nicking DNA via the endonuclease activity of Mre11. This nicking can initiate DSB end resection and HR-mediated reactions (Fig. 7A, right). When DSBs are introduced across the genome via the restriction enzyme AsiSI, some DSBs that are made in transcriptionally active regions are enriched with the HR protein Rad51 (Aymard et al., 2014). We suggest that transcription leading to arrested replication fork arrest recruits the MRX complex and involves DSB end resection.

Interestingly, bidirectional transcription at the E-pro locus is important for rDNA copy number changes, as replacement of an E-pro with a unidirectional *GAL7* promoter in either direction was not sufficient for inducing rDNA copy number changes (Kobayashi and Ganley, 2005). Our findings demonstrated that right-sided transcription induces DSB end resection to initiate HR, but for HR reactions to result in rDNA copy number changes, cohesin associations need to be inhibited by left-sided transcription (Kobayashi *et al*., 2004; Kobayashi and Ganley, 2005). The transcription of noncoding RNA from E-pro is silenced by Sir2, which is recruited to the RFB in a Fob1-dependent manner (Huang and Moazed, 2003). When rDNA stability needs to be maintained, Fob1 recruits Sir2 to E-pro to restrict transcription at E-pro to restrict DSB end resection and not to disrupt cohesin binding. However, when rDNA copy number changes are needed, for example, cells lower *SIR2* expression to activate transcription from E-pro, which stimulates DSB formation and initiates DSB end resection, and HR reactions that are prone to rDNA copy number changes due to the absence of cohesin associations. Because replication forks can stall across the genome, cells may have evolved mechanisms to sense replication stress and inhibit nearby transcription to ensure genome integrity.

## AUTHOR CONTRIBUTIONS

M.S. and T.K. conceived the research and designed the experiments. M.S. performed all the experiments. M.S. wrote the manuscript with input from T.K.

## ACKNOWLEDGEMENTS

We thank members of the Kobayashi laboratory for discussions. We thank Etsuko Ogawara, Machiko Tsumura and Emi Imatani for their technical assistance. This work was supported by Grants-in-Aid for Scientific Research (20H05382 and 18H04709 to M.S. and 17H01443 and 21H04761 to T.K.), the JST FOREST Program (grant number JPMJFR214P to M.S.), JST CREST (grant number JPMJCR19S3 to T.K.), JST AMED (JP21gm1110010 to T.K.) and the Uehara Memorial Foundation, Takeda Science Foundation, and Naito Foundation to M.S.

## STAR METHODS RECOURCE AVAILABILITY

### Lead contact

Further information and requests for resources and reagents should be directed to Mariko Sasaki.

### Materials availability

All the materials used in this study are available upon request to lead contact.

### Data availability

The original data analyzed in this study will be provided upon request to the lead contact.

## EXPERIMENTAL MODELS AND SUBJECTS DETAILS

All yeast strains used in this study are listed in Table S1.

## METHOD DETAILS

### Yeast strains and culture methods for PFGE, rDNA copy number assays, and ERC assays

The yeast strains used in this study are derived from W303 (*MATa ade2-1 ura3-1 his3-11, 15 trp1-1 leu2-3, 112 can1-100 RAD5*) and are listed in **Table S1**. Gene replacement was performed by a one-step procedure using standard methods. Heterozygous gene mutations were introduced into the diploid strain by replacing one allele of the gene of interest. Correct gene replacement was confirmed by PCR-based genotyping. As shown in Fig. 2D and Fig. 3, diploid cells were sporulated by incubating cells in 1% potassium acetate at 30°C for 1–3 days, after which the tetrads were dissected to isolate haploid clones for further analysis. The strains were grown overnight in YP media (1% yeast extract and 2% peptone) supplemented with 2% glucose at 30°C, unless indicated otherwise.

### Culture methods for time course experiments

Time course experiments were performed as described previously (Sasaki and Kobayashi, 2017; 2021). The *GAL-FOB1* strains were grown overnight in YP media (1% yeast extract and 2% peptone) supplemented with 2% glucose at 30°C. The cells were inoculated into ∼200–400 ml of YP media supplemented with 2% raffinose and one drop of antifoam 204 (Sigma) at an appropriate density to reach ∼1 × 10^7^ cells/ml the following morning and were subsequently grown overnight at 23°C. To arrest the cells in the G1 phase of the cell cycle, α-factor (ZYMO RESEARCH) was added to the cultures at a final concentration of 20 nM. The cells were incubated at 23°C for 3–4 hours until >80% of the cells were arrested in G1. For the DSB assays shown in Figs. 1 and 2, 20% galactose was added at a final concentration of 2% to induce *FOB1* expression (*FOB1* ON). The cultures were further incubated for 30 min at 23°C. The cells were collected by centrifugation for 5 min at 4°C at 4,500 × *g*, washed twice with YP medium containing 2% galactose, and resuspended at ∼1 × 10^7^ cells/ml in fresh YP medium containing 2% galactose, 0.075 mg/ml pronase E (Sigma) and one drop of antifoam 204 (Sigma). The cells were incubated at 23°C and collected at various time points.

For the DSB assays shown in Fig. 5B, 5H; ChIP experiments shown in Fig. 4B, 4I, 5I; and Northern blotting shown in Fig. 6, the raffinose-containing preculture where the cells were treated with α-factors was split into two flasks. In one culture, 20% galactose was added at a final concentration of 2% to induce *FOB1* expression (*FOB1* ON). The same volume of 20% raffinose was added to the other cultures (*FOB1* OFF). The cultures were incubated for 30 min at 23°C. The cells were collected, washed, and released into fresh media containing galactose or raffinose at a final concentration of 2%, 0.075 mg/ml pronase E (Sigma) and one drop of antifoam 204 (Sigma). The cells were incubated at 23°C and collected at various time points. For the ChIP experiments shown in Fig. 4B and 4I and Northern blotting shown in Fig. 6, nocodazole was added to a final concentration of 150 g/ml at approximately 30 min after being released from G1 arrest, which arrested cells in the G2/M phase and prevented cells from entering the next cell cycle.

For the northern blotting shown in Fig. 4E and 4G and for the DSB assay shown in Fig. 5E, the raffinose-containing preculture in which the cells were treated with α-factors was split into three flasks. To two of them, 20% galactose was added at a final concentration of 2% to induce *FOB1* expression (*FOB1* ON). After incubating for 30 min at 23°C, the cells were collected and washed. Cells from one culture were released into fresh media that contained galactose at a final concentration of 2%, 0.075 mg/ml pronase E (Sigma) and one drop of antifoam 204 (Sigma). Cells from the other culture were released into fresh media supplemented with galactose and glucose at a final concentration of 2%, 0.075 mg/ml pronase E (Sigma) and one drop of antifoam 204 (Sigma), during which *de novo* transcription was stopped. The last culture was kept in raffinose-containing media throughout the time course. The cells were incubated at 23°C and collected at various time points.

### Genomic DNA preparation

For the PFGE, ERC, and DSB assays, genomic DNA was prepared in low melting temperature agarose plugs as described previously (Sasaki and Kobayashi, 2017; 2021). For the PFGE and ERC assays, cells were collected and washed once with 50 mM EDTA (pH 7.5) (5 × 10^7^ cells per plug). For DSB assays, cells were collected, immediately treated with 0.1% sodium azide, and washed twice with 50 mM EDTA (pH 7.5) (5 × 10^7^ cells per plug). The cells were resuspended in 50 mM EDTA (pH 7.5) (33 μl per plug) and incubated at 42°C. The cell suspension was mixed with a mixture (66 μl per plug) containing 0.83% low-melting-point agarose (SeaPlaque GTG; Lonza), 170 mM sorbitol, 17 mM sodium citrate, 10 mM EDTA (pH 7.5), 0.85% β-mercaptoethanol, and 0.17 mg/ml Zymolyase 100 T (Nacalai) and poured into a plug mold (Bio-Rad). Agarose was allowed to solidify at 4°C for >30 min. Plugs were incubated for 1–1.25 hours at 37°C in a mixture of 450 mM EDTA (pH 7.5), 10 mM Tris-HCl (pH 7.5), 7.5% β-mercaptoethanol and 10 μg/ml RNaseA (Macherey-Nagel). The plugs were then incubated overnight at 50°C in a solution containing 250 mM EDTA (pH 7.5), 10 mM Tris-HCl (pH 7.5), 1% sodium dodecyl sulfate and 1 mg/ml proteinase K (Nacalai). The plugs were washed four times with 50 mM EDTA (pH 7.5) and stored at 4°C in 50 mM EDTA (pH 7.5).

### DSB assays

DSB assays were performed as described previously (Sasaki and Kobayashi, 2017; 2021). Approximately one-third of an agarose plug was cut, transferred to 2.0-ml flat bottom tubes, and equilibrated four times in 1 ml of TE for 15 min at room temperature. Plugs were then equilibrated twice for 30 min at room temperature in 1 ml of 1x NEBuffer 3.1 (New England Biolabs). After the buffer was completely discarded, each plug was incubated in 160 μl of 1x NEBuffer 3.1 containing 1 unit/μl of *Bgl* II (New England Biolabs) overnight at 37°C. Plugs were placed on the tooth of the comb. After extra buffer was removed using lab wipers, the comb was set in the gel tray (15 × 25 cm), and 0.7% agarose in 1x TBE was poured. DNA was separated in 1.45 L of 1x TBE at 2.0 V/cm for 20–22 hours with buffer circulation in a Sub-cell GT electrophoresis system (Bio-Rad). DNA was vacuum-transferred to Hybond-XL (GE Healthcare) as described previously (Sasaki and Kobayashi, 2017; 2021) using VacuGene XL (GE Healthcare). DNA was vacuum-transferred to the membrane at 55 cm Hg for 20 min with 0.25 N HCl, 20 min with denaturing buffer (1.5 M NaCl, 0.5 N NaOH) and 2 hours with transfer buffer (1.5 M NaCl, 0.25 N NaOH). The DNA was fixed to the membrane by soaking it in freshly prepared 0.4 N NaOH for 10 min, followed by washing it with 2x SSC (0.3 M NaCl, 0.03 M citrate) for 10 min.

### PFGE analysis

PFGE analyses were performed as described previously (Sasaki and Kobayashi, 2017; 2021). One-third of a plug and *Hansenula wingei* chromosomal DNA markers (Bio-Rad) were cut and placed on the tooth of the comb. After extra buffer was removed using lab wipers, the comb was set in the gel tray, and 1.0% agarose solution (Pulsed Field Certified Agarose, Bio-Rad) in 0.5x TBE (44.5 mM Tris base, 44.5 mM boric acid and 1 mM EDTA, pH 8.0) was added. PFGE was performed on a Bio-Rad CHEF DR-III system in 2.2 L of 0.5x TBE at 14°C under the following conditions: 68 hours at 3.0 V/cm, 120° included angle, initial switch time of 300 sec, and final switch time of 900 sec. DNA was stained with 0.5 μg/mL ethidium bromide (EtBr) and photographed.

### ERC assays

ERC assays were performed as described previously (Sasaki and Kobayashi, 2021). One-half of an agarose plug was cut and placed on the tooth of the comb. After the extra buffer was removed using lab wipers, the comb was set in a 15 (W) cm × 25 (H) cm gel tray, and 300 ml of 0.4% agarose solution in 1x TBE that was preequilibrated to 50°C was added. After the gel was solidified, the gel was set in a Sub-cell GT Cell System (Bio-Rad), and 1.45 L of 1× Tris-acetate-EDTA (40 mM Tris base, 20 mM acetic acid, and 1 mM EDTA, pH 8.0) was added. Five hundred nanograms of Lambda HindIII DNA markers was loaded, and the DNA was separated by electrophoresis at 1.0 V/cm for ∼72 hr at 4°C with buffer circulation. The buffer was changed every ∼24 hr. DNA was vacuum-transferred to the membrane at 55 cm Hg for 20 min with 0.25 N HCl, 20 min with denaturing buffer (1.5 M NaCl, 0.5 N NaOH) and 2 h with transfer buffer (1.5 M NaCl, 0.25 N NaOH) using a VacuGene XL (GE Healthcare). The DNA was fixed to the membrane by soaking it in freshly prepared 0.4 N NaOH for 10 min, followed by washing it with 2X SSC (0.3 M NaCl, 0.03 M citrate) for 10 min.

### Southern blotting

Southern blotting was performed as described previously (Sasaki and Kobayashi, 2017; 2021).

#### Probe preparation and hybridization

The sequences of primers used to prepare the probes were as follows: probe 1 (5′-ACGAACGACAAGCCTACTCG and 5′-AAAAGGTGCGGAAATGGCTG); probe 2 (5′-CATTTCCTATAGTTAACAGGACATGCC and 5′-AATTCGCACTATCCAGCTGCACTC); and probe 5 (5′-ATGTCGAAAGCTACATATAAGGAAC and 5′-ATGTCAGATCCTGTAGAGACC). To prepare double-stranded DNA probes, PCR products were generated by two rounds of PCR. In brief, the first round of PCR was performed with primers using genomic DNA as a template. The PCR products were gel-purified and used to seed a second round of PCR with the same primers, after which the PCR products were gel-purified. Radioactive probes were generated using the Random Primer DNA Labeling Kit Ver. 2 (TaKaRa) according to the manufacturer’s instructions. Briefly, 50 ng of heat-denatured DNA was incubated for 30 min at 37°C in a total volume of 25 μl containing random primer and 5 μl of [α-^32^P]-dCTP (3,000 Ci/mmol, 10 mCi/ml; Perkin Elmer). After the reaction, 25 μl of probe buffer was added, and unincorporated nucleotides were removed using ProbeQuant G-50 Micro Columns (GE Healthcare) according to the manufacturer’s instructions.

The membrane was prewetted with 0.5 M phosphate buffer (pH 7.2) and prehybridized for >1 hour at 65°C with 25 ml of hybridization buffer (1% bovine serum albumin (Nacalai), 0.5 M phosphate buffer (pH 7.2), 7% sodium dodecyl sulfate, and 1 mM EDTA (pH 8.0)). The membrane was incubated overnight at 65°C with 25 ml of fresh hybridization buffer and a heat-denatured, radiolabeled probe. The membrane was washed four times for 15 min at 65°C with wash buffer (40 mM phosphate buffer [pH 7.2], 1% SDS, and 1 mM EDTA [pH 8.0]) prewarmed at 42°C. The membrane was subsequently exposed to a phosphor screen.

#### Signal quantification

The radioactive signal was detected using a Typhoon FLA7000 (GE Healthcare) and quantified using FUJIFILM Multi Gauge version 2.0 software (Fujifilm). In DSB assays of *GAL-FOB1* strains, the membranes were exposed to phosphor screens for an appropriate exposure time without any saturation of signals. Scanned images were quantified by creating profile regions for all of the lanes and selecting the bands corresponding to the converging forks, arrested forks, linear molecules and DSBs at the same time as setting polygonal background lines for each band. Next, the levels of DSBs and replication intermediates at given time points were calculated as a percentage of the signal intensity of each band relative to the total lane signal, after which the values at t=0 min were subtracted from those at other time points. The DSB frequency was determined by calculating the proportion of the maximum value of the DSB signal to the maximum value of the arrested fork signal in each time course experiment. The same membrane was exposed to phosphor screens for 1 day or longer to increase the signal intensities of DSBs and resected DSBs. Phosphor screens were scanned. The signal intensities of terminal fragments, DSBs and resected DSBs were quantified as described above. The signal intensities of DSBs and resected DSBs were normalized to those of the terminal fragment, and the value at t=0 min was subtracted from the values at other time points. The DSB resection frequency was determined by calculating the ratio of the maximum value of the resected DSB signal to the maximum value of the DSB signal in each time course experiment.

In DSB assays in Gal-pro strains, the membranes were exposed to phosphor screens for 1 day or longer so that signals derived from linear molecules, but not from other molecules, reached saturation. Scanned images were quantified by creating profile regions for all of the lanes and selecting the bands corresponding to arrested forks, DSBs, resected DSBs, and the ∼4.3 kb fragment that contains the restriction fragment around the endogenous *URA3* at the same time. Because the resected DSB signal was very low and spread over a wide region, we constructed straight background lines for each band. The signal intensities of arrested forks, DSBs, and resected DSBs were normalized to those of the endogenous *URA3* fragment, and the value at t=0 min was subtracted from the values at other time points. The DSB frequency was determined by calculating the proportion of the maximum value of the DSB signal to the maximum value of the arrested fork signal in each time course experiment. The DSB resection frequency was determined by calculating the ratio of the maximum value of the resected DSB signal to the maximum value of the DSB signal in each time course experiment.

For ERC detection, the membranes were exposed to a phosphor screen for an appropriate exposure time without any saturation of signals. The radioactive signal was detected using a Typhoon FLA7000 (GE Healthcare), and the scanned image was used to quantitate the genomic rDNA signal. The membranes were exposed to a phosphor screen for a few days, and the scanned image was used to quantify the ERCs. Using FUJIFILM Multi Gauge version 2.0 software (Fujifilm), scanned images were quantified by creating profile regions for all of the lanes and selecting the bands of genomic rDNA or ERCs at the same time as setting polygonal background lines for each band. The frequency of ERCs was determined by calculating the ratio of the ERC signal to the genomic rDNA signal in each lane and was compared between different strains.

### Chromatin immunoprecipitation-quantitative PCR analysis

Thirty-five millilitres of cultures from the time course experiments (3.5 × 10^8^ cells) was transferred to 50 ml conical tubes, to which 972 μl of 37% formaldehyde was added at a final concentration of 1%, and the cells were crosslinked by shaking for 30 min at room temperature on a reciprocal shaker at 60 r/min. The crosslinking reaction was quenched by adding glycine to a final concentration of 125 mM and shaking for 5 min at room temperature on a reciprocal shaker at 60 r/min. The cells were collected by centrifugation at 2,380 × *g* for 3 min at 4°C, washed twice with 20 ml of TBS (20 mM Tris, 150 mM NaCl, pH 7.4), and washed once with 5 ml of FA lysis buffer (50 mM HEPES-KOH, pH 7.4; 150 mM NaCl; 1 mM EDTA, pH 8.0; 1% Triton X-100; 0.1% sodium deoxycholate; 0.1% SDS). The cell pellets were resuspended in 0.5 ml of FA lysis buffer, transferred to a 1.5 ml microtube, and collected by centrifugation at 2,380 × *g* for 1 min at 4°C.

The cells were resuspended in 400 μl of FA lysis buffer supplemented with 1 mM PMSF and 1x Complete EDTA-free solution and transferred to a 2 ml tube (STC-0250, Yasui Kikai). Zirconia/silica beads (1.5 g) 0.5 mm in diameter were added to each tube. The cells were disrupted using a Multibead Shocker (Yasui Kikai) for 20 cycles of 30 sec on at 2,700 rpm and 30 sec off at 2°C. A burned 23–26-G needle was used to make a few holes at the bottom of the tube, which was inserted into a 5 ml tube and centrifuged at 1,000 × *g* for 30 sec at 4°C. The cell lysate was transferred to a 1.5 ml TPX tube. Chromatin was sheared by sonication at 4°C at high power with ice on a Bioruptor for three sets of 5 cycles of 30 sec on and 30 sec off. Sheared chromatin samples were centrifuged at 20,380 *g* for 15 min at 4°C and recovered with the same volume of FA lysis buffer. A total of 10 μl was set aside as input DNA.

A total of 320 μl was transferred to a low-binding tube (Costar) and incubated with 10 μl of Dynabeads Protein G (Thermo Fisher) for 1 hr at 4°C, after which the precleared samples were transferred to a new low-binding tube. Then, 2.1 μg of anti-Flag antibody (M2; Sigma‒ Aldrich) was added to each tube, and the mixture was incubated for 3 hrs at 4°C. The chromatin samples were incubated with an antibody and then incubated overnight at 4°C with 12 μl of Dynabeads Protein G, which was preincubated with 5 mg/ml BSA solution in PBS. The beads were washed twice with 0.5 ml of FA lysis buffer, twice with 0.5 ml of FA lysis buffer supplemented with 360 mM NaCl, twice with 0.5 ml of ChIP wash buffer (10 mM Tris-HCl (pH 8.0), 250 mM lithium chloride, 1 mM EDTA (pH 8.0), 0.5% NP-40 and 0.5% sodium deoxycholate), and twice with 0.5 ml of 1x TE buffer. After the buffer was completely removed, 54 μl of ChIP elution buffer (50 mM Tris-HCl (pH 8.0), 10 mM EDTA, and 1% SDS) was added, and the immunoprecipitates were eluted from the beads by incubating for 15 min at 65°C. Fifty μl of eluate was transferred to a new tube and mixed with 150 μl of TE containing 1% SDS. Ten μl of input sample was mixed with 190 μl of TE containing 1%. To each sample, RNaseA was added at a final concentration of 50 μg/ml. These samples were incubated overnight at 65°C to reverse crosslinking and digest RNA. Then, proteinase K was added at a final concentration of 0.75 mg/ml, and the mixture was incubated for 2 hr at 50°C. DNA was purified using a QIAquick PCR Purification Kit (QIAGEN). DNA was eluted with 100 μl of dH_2_O.

qPCR was performed on a Thermal Cycler Dice Real Time System II (TaKaRa) with THUNDERBIRD SYBR qPCR Mix or Next SYBR qPCR Mix (TOYOBO) according to the manufacturers’ instructions. One of the input samples was diluted serially to 1/20, 1/200, 1/2,000 and 1/20,000 dilutions, which were subsequently used to construct the standard curve. All the input samples were subsequently diluted 400-fold. IP samples were diluted 10-fold. qPCR was performed with all the diluted samples as well as the standard. The amount of DNA in each sample was subsequently determined relative to the standard. After considering dilution factors, % input was expressed for each time point of each culture as the amount of DNA in the IP relative to that in the input.

### Isolation of total RNA

To isolate RNA, 2 × 10^8^ cells were collected. The cells were resuspended in 1 ml of ice-cold dH_2_O, divided into two 1.5 ml tubes, and collected by centrifugation at 2,380 *g* for 1 min at 4°C. The cells were washed with 0.5 ml of ice-cold dH_2_O and centrifuged at 2,380 × *g* for 1 min at 4°C. The cells were resuspended in 400 μl of TES [10 mM Tris-HCl (pH 7.5), 10 mM EDTA (pH 7.5), and 0.5% SDS]. To the cell suspension, 400 μl of acid-phenol:chloroform (pH 4.5) (Thermo Fisher Scientific) was added, and the mixture was vortexed. The cell suspensions were incubated for 1 hr at 65°C, during which the samples were mixed by vortexing every 15 min. After the samples were chilled on ice for 5 min, they were centrifuged for 10 min at 4°C at 20,380 × *g*. The supernatant (<400 μl) was transferred to a new tube. To each tube, 400 μl of acid-phenol:chloroform (pH 4.5) was added, and the solution was mixed by vortexing. The samples were centrifuged for 10 min at 4°C at 20,380 × *g*. From two tubes per sample, 200 μl of supernatant was transferred to a single, new tube (400 μl in total per sample).

RNA was precipitated by adding 0.1 volume of 3 M NaOAc (pH 5.2) and 2.5 volume of 100% ethanol to the tube, mixing the mixture by inverting the tube a few times, and storing the mixture overnight at –20°C. RNA was collected by centrifugation for 15 min at 4°C and 20,380 *g*. The supernatant was removed by aspiration. To wash the RNA pellets, 400 μl of ice-cold 70% ethanol was added, the tube was centrifuged for 30 sec at 4°C and 20,380 × *g,* and the ethanol was removed by aspiration. The RNA was resuspended in diethylpyrocarbonate (DEPC)-treated dH_2_O. The concentration of RNA was determined by measuring the OD_260_ on a Nanodrop. The RNA was stored at –80°C.

### Agarose gel electrophoresis of RNA

Each sample was separated in two agarose gels, one for detection of IGS1-F and the other for IGS1-R and IGS2-R. Thirty micrograms of total RNA was added to 7 μl of DEPC-treated dH_2_O and mixed with 17 μl of RNA sample buffer (396 μl of deionized formamide and 120 μl of 10x MOPS buffer [0.2 M MOPS, 50 mM sodium acetate pH 5.2, 10 mM EDTA pH 7.5 in DEPC-treated dH_2_O], and 162 μl of 37% formaldehyde). To prepare the RNA size markers for loading into two gels, 3.6 μg of DynaMarker RNA High markers was added to 4.8 μl of DEPC-treated dH_2_O, which was subsequently mixed with 11.2 μl of RNA sample buffer. The samples were heated at 65°C for 20 min, followed by rapid chilling on ice. Six microliters of 6x gel loading dye (B7025S, New England Biolabs) and 1.5 μl of 2.5 mg/ml EtBr were added to each RNA sample. 4 μl of 6x Gel Loading Dye (B7025S, New England Biolabs) and 1 μl of 2.5 mg/ml EtBr were added to the RNA size markers.

Two 1% agarose gels were prepared in a 15 cm (W) × 10 cm (H) gel tray (100 ml agarose solution per gel). One gram of agarose powder was dissolved in 73 ml of DEPC-treated dH_2_O by microwave. After the solution was cooled to 60°C, 17 ml of 37% formaldehyde and 10 ml of 10x MOPS buffer were added and mixed gently. The solution was poured into a gel tray, allowed to solidify, and placed in a Wide Mini-Sub Cell GT System (Bio-Rad). A total of 650 ml of 1x MOPS buffer in DEPC-treated dH_2_O was poured into the electrophoresis tank. The gels were prerun at 50 V (∼3.4 V/cm) for 10 min. The wells were thoroughly washed. Ten microliters (10 μg of total RNA) of RNA sample and of RNA size markers were loaded into each gel. The gels were run at 20 V (∼1.4 V/cm) for 30 min and then at 60 V (∼4 V/cm) until the pink loading dye was reached in 6x gel loading dye (B7025S, New England Biolabs), which shows a similar migration speed to that of bromophenol blue, which migrated approximately 2/3 of the gel. After electrophoresis, the gel was rinsed for 10 min with ∼200 ml of DEPC-treated dH_2_O and photographed. RNA was vacuum-transferred to Hybond-N+ (GE Healthcare) at 55 cm Hg for 1.5 hrs with RNA transfer buffer (3 M NaCl, 0.01 N NaOH) using a VacuGene XL (GE Healthcare). The membrane was rinsed with 2x SSC for 10 min. RNA was fixed to the membrane by UV crosslinking at 120 mJ/cm^2^.

### Northern blotting

To prepare strand-specific probes 3 and 4, double-stranded DNA fragments were first amplified via PCR using the following primers and genomic DNA as a template: probe 3 (5′-TCGCCAACCATTCCATATCT and 5′-CGATGAGGATGATAGTGTGTAAGA) and probe 4 (5′-AGGGAAATGGAGGGAAGAGA and 5′-TCTTGGCTTCCTATGCTAAATCC). The PCR products were gel-purified and used to seed a second round of PCR with the same primers, after which the PCR products were gel-purified. Single-stranded, radiolabeled probes were generated by linear PCR. The reactions were set up in a final volume of 20 μl containing 50 ng of gel-purified PCR products as a template, 0.2 mM dATP, 0.2 mM dTTP, 0.2 mM dGTP, 5 μl of [α-^32^P]-dCTP (3,000 Ci/mmol, 10 mCi/ml; Perkin Elmer), 0.5 μM primer, 1.25 units of ExTaq (TaKaRa) and 1x ExTaq buffer. The primers used were 5′-AGTTCCAGAGAGGCAGCGTA for probe 3 and 5′-CATTATGCTCATTGGGTTGC for probe 4. PCR was initiated by a denaturation step at 94°C for 2 min, followed by 35 cycles of amplification (94°C for 20 sec, 51°C for 20 sec, and 72°C for 30 sec) and a final step at 72°C for 2 min. After the reaction, 30 μl of probe buffer was added, and unincorporated nucleotides were removed using ProbeQuant G-50 Micro Columns (GE Healthcare) according to the manufacturer’s instructions.

The membrane was prewetted with 2x SSC and incubated for >1 hour at 42°C with 10 ml of ULTRAhyb ultrasensitive hybridization buffer (Thermo Fisher). The radiolabeled probe prepared by linear PCR as above was heat denatured for 5 min at 100°C and immediately added to the hybridization buffer. The membrane was incubated overnight at 42°C. The membrane was rinsed twice with 2x SSC and washed twice for 15 min at 42°C with Northern wash buffer 1 (2x SSC, 0.1x SDS) prewarmed at 42°C. The membrane was washed twice for 15 min at 42°C with Northern wash buffer 2 (0.1x SSC, 0.1x SDS) prewarmed at 42°C. The membrane was subsequently exposed to a phosphor screen. The radioactive signal was detected using a Typhoon FLA7000 (GE Healthcare). The signal was quantitated by FUJIFILM Multi Gauge version 2.0 software (Fujifilm).

To strip the probes, the membrane was placed in tupperware and incubated with ∼200 ml of boiling 0.1% SDS solution for 30 min by shaking on a reciprocal shaker, followed by washing with 2x SSC. To prepare the TSA1 probe for use as an RNA loading control, a double-stranded probe was prepared by two rounds of PCR as described above using primers (5′-CAAGTTCAAAAGCAAGCTCC and 5′-ACCAATCTCAAGGCTTCGTC). Radioactive probes were generated using the Random Primer DNA Labeling Kit Ver. 2 (TaKaRa) and purified using ProbeQuant G-50 Micro Columns (GE Healthcare) as described above. The membrane was hybridized to the TSA1 probe in ULTRAhyb Ultrasensitive Hybridization Buffer (Thermo Fisher), washed, exposed to a phosphor screen, and quantitated as described above.

## QUANTIFICATION AND STATISTICAL ANALYSIS

Statistical analysis was performed using GraphPad Prism software (version 6.0d or 6.0h). One-way ANOVA was used to compare differences among three or more groups. Tukey’s multiple comparisons tests were performed to identify statistically significant differences between different groups. The two-sided Student t-test was used to compare difference between two groups The statistical parameters used are described in the figures, figure legends and METHODS section.

