## Supplemental Table 1 for "Transcription near arrested DNA replication forks triggers ribosomal DNA copy number changes": 20231216-Supplementary Table 1.pdf

**Supplementary Table 1. *S. cerevisiae* strains used in this study**

| Name | Genotype |
|---|---|
| MSY360 | *MATa, NatNT2-GALL-FOB1, bar1::LEU2* |
| MSY375 | *MATa, NatNT2-GALL-FOB1, bar1::LEU2, ctf4Δ::kanMX* |
| MSY937 | *MATa, NatNT2-GALL-FOB1, bar1::LEU2, sir2Δ::hphMX, hmlΔ::kanMX* |
| MSY967 | *MATa, NatNT2-GALL-FOB1, bar1::LEU2, sir2Δ::hphMX, hmlΔ::kanMX, ctf4Δ::TRP1* |
| MSY1585 | *MATa, NatNT2-GALL-FOB1, bar1::LEU2, sir2Δ::hphMX, hmlΔ::kanMX, mre11Δ::klTRP1* |
| MSY1027 | *MATa, NatNT2-GALL-FOB1, bar1::LEU2, sir2Δ::hphMX, hmlΔ::kanMX, sae2Δ::klTRP1* |
| MSY1023 | *MATa, NatNT2-GALL-FOB1, bar1::LEU2, sir2Δ::hphMX, hmlΔ::kanMX, rad52Δ::klTRP1* |
| MSY1583 | *MATa/α, fob1::LEU2/FOB1, sir2Δ::hphMX/SIR2, mre11Δ::klTRP1/MRE11* |
| MSY769 | *MATa/α, fob1::LEU2/FOB1, rad52Δ::hphMX/RAD52, sir2Δ::kanMX/SIR2* |
| MSY476 | *MATa, NatNT2-GALL-FOB1, bar1::LEU2, MCD1-6His10FLAG::kanMX* |
| MSY1459 | *MATa, NatNT2-GALL-FOB1, bar1::LEU2, MCD1-6His10FLAG::kanMX, sir2Δ::hphMX, hmlΔ::HIS3MX* |
| MSY988[a] | *MATa, NatNT2-GALL-FOB1, bar1::LEU2, E-proΔ::GAL1/10p-URA3* |
| MSY1456 | *MATa, NatNT2-GALL-FOB1, bar1::LEU2, E-proΔ::GAL1/10p-URA3, MCD1-6His10FLAG::kanMX* |
| MSY1535 | *MATa, NatNT2-GALL-FOB1, bar1::LEU2, E-proΔ::GAL1/10p-URA3, rad50Δ::kanMX* |
| MSY1541 | *MATa, NatNT2-GALL-FOB1, bar1::LEU2, E-proΔ::GAL1/10p-URA3, sae2Δ::kanMX* |
| MSY1543 | *MATa, NatNT2-GALL-FOB1, bar1::LEU2, E-proΔ::GAL1/10p-URA3, rad52Δ::hphMX* |

All strains are derivatives of W303, which is *ade2-1, ura3-1, his3-11, 15, trp1-1, leu2-3, 112, can1-100*, and *RAD5*.

[a]This strain was constructed from TAK2004a that carries ~80 rDNA copies and in which E-pro is replaced by the *GAL1/10* promoter.
